# Chromosome-level assembly of the Atlantic silverside genome reveals extreme levels of sequence diversity and structural genetic variation

**DOI:** 10.1101/2020.10.27.357293

**Authors:** Anna Tigano, Arne Jacobs, Aryn P. Wilder, Ankita Nand, Ye Zhan, Job Dekker, Nina O. Therkildsen

## Abstract

The levels and distribution of standing genetic variation in a genome can provide a wealth of insights about the adaptive potential, demographic history, and genome structure of a population or species. As structural variants are increasingly associated with traits important for adaptation and speciation, investigating both sequence and structural variation is essential for wholly tapping this potential. Using a combination of shotgun sequencing, 10X Genomics linked reads and proximity-ligation data (Chicago and Hi-C), we produced and annotated a chromosome-level genome assembly for the Atlantic silverside (*Menidia menidia*) - an established ecological model for studying the phenotypic effects of natural and artificial selection - and examined patterns of genomic variation across two individuals sampled from different populations with divergent local adaptations. Levels of diversity varied substantially across each chromosome, consistently being highly elevated near the ends (presumably near telomeric regions) and dipping to near zero around putative centromeres. Overall, our estimate of the genome-wide average heterozygosity in the Atlantic silverside is the highest reported for a fish, or any vertebrate, to date (1.32-1.76% depending on inference method and sample). Furthermore, we also found extreme levels of structural variation, affecting ~23% of the total genome sequence, including multiple large inversions (> 1 Mb and up to 12.6 Mb) associated with previously identified haploblocks showing strong differentiation between locally adapted populations. These extreme levels of standing genetic variation are likely associated with large effective population sizes and may help explain the remarkable adaptive divergence among populations of the Atlantic silverside.

## Introduction

Standing genetic variation is widely recognized as the main source of adaptation (Barrett & Schluter 2008; Tigano & Friesen 2016) and is important for natural populations to maximize their potential to adapt to changes in their environment. As genetic diversity is the result of the interplay of mutation, selection, drift and gene flow, the levels and patterns of standing genetic variation found within a species can provide important insights not only about its adaptive potential but also about its demographic and evolutionary history.

Traditionally, quantification of standing genetic variation has been based on sequence variation, often across a limited number of genetic markers, or small microsatellite repeats. As an increasing number of empirical studies shows the mosaic nature of the genome (Pääbo 2003) with different genomic regions showing vastly different levels of diversity and differentiation (e.g., Martinez Barrio et al. 2016; Campagna et al. 2017; Murray et al. 2017; Sardell et al. 2018), it is evident that small marker panels do not grant the resolution to describe variation in diversity across the genome (Dutoit et al. 2016). Furthermore, structural variation, including changes in the position, orientation, and number of copies of DNA sequence, is generally neglected as a type of standing genetic variation. Structural variation has been associated directly or indirectly with many traits involved in speciation and adaptation and is abundant in the few genomes in which they have been catalogued (Wellenreuther & Bernatchez 2018; Catanach et al. 2019; Lucek et al. 2019; Mérot et al. 2020; Tigano et al. 2020; Weissensteiner et al. 2020). Structural variants can directly affect phenotypic traits, such as the insertion of a repeated transposable element in the iconic case of industrial melanism in the peppered moth (*Biston betularia*; Van’t Hof et al. 2016), or may promote the maintenance of divergent haplotypes between locally adapted populations or groups (e.g. ecotypes or morphs) within single populations via recombination suppression (e.g., Faria et al. 2019; Kess et al. 2020). Structural variation is therefore a key source of standing genetic variation, which can also play an important role in rapid evolutionary responses to environmental change (Reid et al. 2016). To better assess levels of standing variation and understand how demographic and evolutionary factors contribute to their distribution in the genome, we need to examine large proportions of the genome, preferably its entirety, and examine sequence and structural variation jointly. A high-quality reference genome for the species of interest is therefore fundamental as we need both broad coverage to accurately assess variation in levels of standing sequence variation across the genome, and high contiguity to investigate standing structural variation.

The Atlantic silverside (*Menidia menidia*), a small coastal fish distributed along the Atlantic coast of North America, shows a remarkable degree of local adaptation in a suite of traits, including growth rate, number of vertebrae, and temperature-dependent sex determination (Hice et al. 2012), that are associated with strong environmental gradients across its wide latitudinal range. This species also provided the first discovery of temperature-dependent sex determination in fishes (Conover & Kynard 1981) and was one of the first species in which countergradient phenotypic variation was documented (Conover & Present 1990). Through extensive prior work, the Atlantic silverside has, in fact, become an important ecological model to study the phenotypic effects of selection, both natural and artificial, in the wild and under controlled conditions in the lab (Conover & Munch 2002; Conover et al. 2005; Hice et al. 2012). In one iconic experiment, wild-caught Atlantic silversides were subjected to different size-selective regimes to investigate the potential of fisheries to induce evolutionary change in harvested species (Conover & Munch 2002). Seventeen years later, genomic analysis of fish from that experiment identified substantial allele frequency shifts associated with rapid phenotypic shifts in growth rates (Therkildsen et al. 2019). In the absence of a reference genome, genomic reads were mapped to the silverside reference transcriptome, so only protein-coding regions of the genome were analyzed (‘in-silico’ exome capture). Yet, anchoring the transcriptome contigs to the medaka (*Oryzias latipes*) chromosome-level reference genome revealed that the most conspicuous allele frequency shifts clustered into a single block on chromosome 24, where more than 9,000 SNPs in strong linkage disequilibrium (LD) increased from low (< 0.05) to high frequency (~0.6) in only five generations. Additional data from natural populations across the geographical distribution of the species showed that this same block, likely spanning several Mb of the chromosome, was fixed for opposite haplotypes among wild silverside populations that naturally differ in growth rates (Conover & Present 1990; Conover & Munch 2002; Therkildsen et al. 2019). Moreover, three additional blocks comprising hundreds of genes in high linkage disequilibrium (LD) were found to be segregating among the natural populations, with each LD block (‘haploblocks’ hereafter) mapping predominantly to unique medaka chromosomes (Wilder et al. 2020). Similar to the haploblock on chromosome 24, opposite haplotypes in these haploblocks were nearly fixed between natural populations that otherwise showed low genome-wide pairwise differentiation. Furthermore, strong LD between genes in these blocks suggested that local recombination suppression, possibly due to inversions, and natural selection maintained these divergent haploblocks in the face of gene flow. It thus appears that large haploblocks play an important role in maintaining local adaptations in the Atlantic silverside, although the exact extent of the genome spanned by these haploblocks and the genomic mechanism maintaining LD are unknown.

Given the wealth of ecological information available for the Atlantic silverside and its potential as an evolutionary model to study adaptation and fishery-induced evolutionary change, developing genomic resources for this species is timely and holds great potential for addressing many pressing questions in evolutionary and conservation biology. Previous population genomic analyses based on the transcriptome reference anchored to the medaka genome were limited to the coding genes and, given the unknown degree of synteny conservation between the Atlantic silverside and the medaka, how variants relevant to adaptation and fishery-induced selection clustered in the genome was uncertain. To enable analysis of both coding and non-coding regions, to accurately estimate levels and the genomic distribution of standing genetic variation, both sequence and structural, and to reconstruct the specific genomic structure of the Atlantic silverside genome, we produced a chromosome-level genome assembly for the species using a combination of genomic approaches. Because of known geographic differentiation, we estimated levels of sequence variation within genomes from both the southern and northern parts of the distribution and characterized standing structural variation between these two genomes. Finally, we tested whether the haploblocks identified on four different chromosomes between southern and northern populations were associated with large inversions as the patterns of differentiation and LD suggested (Therkildsen et al. 2019). Our work illustrates the wealth of information that can be obtained from the analysis of one or two genomes in the presence of a high quality reference sequence, and shows that, to the best of our knowledge, the Atlantic silverside has the highest nucleotide diversity reported for a vertebrate to date, and extreme levels of structural variation between two locally adapted populations. The distribution of diversity across the genome is strongly affected by structural variants and, seemingly, by genome features such as centromeres and telomeres. These results taken together highlight the importance of high-quality genomic resources as they enable the joint analysis of sequence and structural variation at the whole-genome level.

## Methods

### Reference genome assembly

We built a reference genome for the Atlantic silverside through three steps: First, we created a draft assembly using 10X Genomics linked-reads technology (10X Genomics, Pleasanton, CA, USA); second, we used proximity ligation data - Chicago® (Putnam et al. 2016) and Dovetail™ Hi-C (Lieberman-Aiden et al. 2009) - from Dovetail Genomics to increase contiguity, break up mis-joins, and orient and join scaffolds into chromosomes; and finally, we used short-insert reads to close gaps in the scaffolded and error-corrected assembly. The data were generated from muscle tissue dissected from two lab-reared F1 offspring of Atlantic silversides collected from the wild on Jekyll Island, Georgia, USA (N 31.02, W 81.43; the southern end of the species distribution range) in May 2017. For 10X Genomics library preparation, we extracted DNA from fresh tissue from one individual using the MagAttract HMW DNA Kit (Qiagen). Prior to library preparation, we selected fragments longer than 30 kb using a BluePippin device (Sage Science). A 10X Genomics library was prepared following standard procedure and sequenced using two lanes of paired-end 150 bp reads on a HiSeq2500 (rapid run mode) at the Biotechnology Resource Center Genomics Facility at Cornell University. To assemble the linked reads, we ran the program *Supernova* (Weisenfeld et al. 2017) from 10X Genomics with varying numbers of reads and compared assembly statistics to identify the number of reads that resulted in the most contiguous assembly. Tissue from the second individual was flash-frozen in liquid nitrogen and shipped to Dovetail Genomics, where Chicago and Hi-C libraries were prepared for further scaffolding. These long-range libraries were sequenced in one lane of Illumina HiSeq X using paired-end 150 bp reads. Two rounds of scaffolding with *HiRise™*, a software pipeline developed specifically for genome scaffolding with Chicago and Hi-C data, were run to scaffold and error-correct the best 10X Genomics draft assembly using Dovetail long-range data. Finally, the barcode-trimmed 10X Genomics reads were used to close gaps between contigs.

For each of the intermediate and the final assemblies we produced genome contiguity and other assembly statistics using the *assemblathon_stats.pl* script from the Korf Laboratory (https://github.com/KorfLab/Assemblathon/blob/master/assemblathon_stats.pl) and assessed assembly completeness with *BUSCO v3* (Simão et al. 2015) using the Actinopterygii gene set (4584 genes).

We estimated the genome size and heterozygosity (i.e. the nucleotide diversity π within a single individual) from the raw 10X Genomics data using a k-mer distribution approach. We removed barcodes with the program *longranger basic*, trimmed all reads to the same length of 128 bp (as read length is in the equation to estimate genome size) with *cutadapt* (Martin 2011), and estimated the distribution of 25-mers using *Jellyfish* (Marçais & Kingsford 2011). Finally, we analyzed the 25-mers distribution with the web application of *GenomeScope* (Vurture et al. 2017), which runs mixture models based on the binomial distributions of k-mer profiles to estimate genome size, heterozygosity and repeat content.

### Synteny with medaka

The chromosome-level genome assembly of medaka (*Oryzias latipes*) was used by Therkildsen et al. (2019) to order and orient contigs of the Atlantic silverside transcriptome (Therkildsen & Baumann 2020). Although the two species carry the same number of chromosomes (Uwa & Ojima 1981; Warkentine et al. 1987) and few interchromosomal rearrangements have been observed between other species within the Atherinomorpha clade (Amores et al. 2014; Miller et al. 2019), the estimated divergence time between medaka and Atlantic silverside is 91 million years (estimate based on 15 studies, timetree.org) and the degree of syntenic conservation between the two species was unknown. We assessed synteny between the two species using the newly assembled Atlantic silverside reference genome. We aligned the silverside genome to the medaka genome (GenBank assembly accession GCA_002234675.1) with the *lastal* program in *LAST* (Kiełbasa et al. 2011; Frith & Kawaguchi 2015) using parameters optimized for distantly related species (*-m100-E0.05*). Given the deep divergence between the two species, we kept low-confidence alignments (*last-split-m1*). We filtered alignments shorter than 500 bp and visualized syntenic relationships only for silverside scaffolds longer than 1 Mb (‘chromosome assembly’, see below) using the software CIRCA (omgenomics.com/circa).

### Repeat and gene annotation

We annotated the Atlantic silverside genome using a combination of the *BRAKER2* (Hoff et al. 2019) and *MAKER* (Holt & Yandell 2011) pipelines, which combine repeat masking, *ab initio* gene predictor models and protein and transcript evidence for *de novo* identification and annotation of genes. To annotate repetitive elements, we first identified repeats *de novo* in the Atlantic silverside genome using *Repeatmodeler* (Smit & Hubley 2008) and NCBI as a search engine and combined the resulting species-specific library with a library of known repeats in teleosts (downloaded from the RepBase website (Bao et al. 2015) in July 2018). The merged libraries were then used to annotate repeats in the Atlantic silverside genome with *Repeatmasker* (Smit et al. 2015). We then filtered annotated repeats to only keep complex repeats for soft-masking. Next, we used *BRAKER2* to train *AUGUSTUS* (Stanke et al. 2006; Stanke et al. 2008; Buchfink et al. 2015) on the soft-masked genome with unpublished mRNA-seq evidence from 24 Atlantic silverside individuals from different populations and developmental stages, along with protein homology evidence from six different teleost species (medaka [*Oryzias latipes*], tilapia [*Oreochromis aureus*], platyfish [*Xiphophorus maculatus*], zebrafish [*Danio rerio*], stickleback [*Gasterosteus aculeatus*] and fugu [*Takifugu rubripes*]), which were downloaded from ensemble.org (Ensembl 98; Cunningham et al. 2019) and the UniProtKB (Swiss-Prot) protein database. Second, we ran five rounds of annotation in *MAKER* using different input datasets. The first round of *MAKER* was performed on the genome with only complex repeats masked using the non-redundant transcriptome of Atlantic silverside (Therkildsen and Palumbi 2017, Therkildsen and Baumann 2020) as mRNA-seq evidence, and the six protein sequence datasets from other species as protein homology evidence. We then trained *SNAP* (Korf 2004) on the output of the initial *MAKER* run for *ab initio* gene model prediction. We ran *MAKER* a second time adding the SNAP *ab initio* gene predictions. Using the *MAKER* output from this second round, we re-trained *SNAP* and ran *MAKER* three additional times (round 3 to 5) including the updated *SNAP* gene predictions, the *AUGUSTUS* gene predictions from *BRAKER2* and the updated *MAKER* annotation.

Lastly, we performed a functional annotation using *Blast2GO* in *Omnibox v.1.2.4* (Götz et al. 2008) utilizing the UniProtKB (Swiss-Prot) database and *InterProScan2* results. Annotated Atlantic silverside nucleotide sequences for all predicted genes were blasted against the UniProtKB database using *DIAMOND* v. 0.9.34 (Buchfink et al. 2015) with an e-value cutoff of 10^-5^. *InterProScan2* was used to annotate proteins with *PFAM* and *Panther* annotations and identify GO terms. *Blast2GO* default mapping and annotation steps were performed using both lines of evidence to create an integrated annotation file.

### Comparison of sequence and structural standing genetic variation between populations

As Atlantic silversides from Georgia show strong genomic differentiation from populations further north, primarily concentrated in large haploblocks on four chromosomes (Therkildsen et al. 2019; Wilder et al. 2020), we also sequenced the genome of a representative individual from Mumford Cove, Connecticut, USA (N 41.32°, W 72.02°) collected in June 2016 for comparison. Genomic DNA was extracted from muscle tissue using the DNeasy Blood and Tissue kit (Qiagen) and normalized to 40 ng/μl. We prepared a genomic DNA library using the TruSeq DNA PCR-free library kit (Illumina) following the manufacturer’s protocol for 550 bp insert libraries. The shotgun library was sequenced using paired-end 150 bp reads on an Illumina HiSeq4000.

We estimated genome size and heterozygosity from the raw data from this shotgun library using the same k-mer approach as for the Georgia individual described above. To compare our heterozygosity estimates in Atlantic silversides from Connecticut and Georgia with other fish species, we searched the literature for heterozygosity estimates from Genomescope with the keywords “Genomescope heterozygosity fish”, or from variant calling methods in other fish genomes, using Google Scholar. We also estimated heterozygosity directly by calculating the proportion of heterozygous sites in each genome. For the Georgia individual we used the processed 10X data as above. For the Connecticut individual we trimmed adapters and low-quality data from the raw shotgun data in *Trimmomatic* (Bolger et al. 2014). We mapped data from the two libraries to the chromosome assembly (only the largest 27 scaffolds-see Results) with *bwa mem* (Li & Durbin 2009) and removed duplicates with *samblaster* (Faust & Hall 2014). We called variants with *bcftools mpileup* and *bcftools call* (Danecek et al. 2014). As areas of the genome covered by more than twice the mean sequencing depth could represent repetitive areas or assembly artefact, we calculated genome coverage for each of the two libraries with *genomeCoverageBed* from *BEDtools* (Quinlan & Hall 2010) and identified the depth mode from the calculated distribution (95x for the southern genome and 74x for the northern genome). We then filtered variants that were flagged as low-quality, that had mapping quality below 20, sequencing depth below 20, and more than twice the mode sequencing depth for each of the two libraries using *bcftools filter* (Li et al. 2009). To accurately estimate the proportion of heterozygous sites in the genome, we subtracted the number of sites that had sequencing depth below 20 and above twice the mode sequencing depth from the total genome size (to get the sum of sites that could be identified as either homozygous or heterozygous based on our criteria). To visualize variation along the genome, we plotted estimates of heterozygosity in 50-kb sliding windows along the genome for each of the two individuals using the *qqman* package (Turner 2014) in R (R Core Team 2019). To assess the reduction in diversity in protein-coding regions due to positive and purifying selection, we calculated heterozygosity in the regions annotated as coding sequences only and compared this to the genome-wide estimate.

Finally, we identified structural variants (SVs) segregating between the Connecticut and Georgia genomes using *Delly2 v.0.8.1* (Rausch et al. 2012). For this analysis we used the shotgun library data (74x coverage) from Connecticut mapped to the Georgia reference genome as described above. We called SVs using the command *delly call* and default settings. As genotyping a single individual in *Delly* is prone to false positives we applied the following stringent filters: We retained only homozygous SVs (*vac=2*) that passed quality filters (*PASS*) and that had at least 20 reads supporting the variant calls, whether they came from paired-end clustering or split-read analysis or a combination of the two, but not more than 100 reads since these could be due to repetitive elements in the genome. As *Delly2* outputted redundant genotypes, e.g. inversions that had slightly different breakpoints were reported as independent variants, we used *bedtools merge* to merge these overlapping features. To validate duplication calls we also calculated coverage for each of these variants and retained only those putative duplications that had coverage more than 1.8-fold the whole genome sequencing depth (74x).

To confirm the large SVs observed between the two genomes examined, we generated a second Hi-C library from an Atlantic silverside individual caught in Mumford Cove, Connecticut in June 2016 (different from the sample used for the shotgun assembly). Liver tissue was excised and digested for 2 hours in collagenase digestion buffer (perfusion buffer plus 12.5 μM CaCl2 plus collagenases II and IV (5 mg/ml each)). The cell suspension was then strained through a 100 μm cell strainer, washed with 1 ml cold PBS three times, resuspended in 45 ml PBS, and quantified in a hemocytometer. The cross-linking protocol was modified from Belton et al. (2012) as follows. 1.25 ml of 37% formaldehyde was added twice to the cell preparation, then incubated at room temperature for 10 minutes, inverting every 1-2 minutes. To quench the formaldehyde in the reaction, 2.5 ml of 2.5 M glycine was added three times. The sample was incubated at room temperature for 5 minutes, then on ice for 15 minutes to stop the cross-linking. The cells were pelleted by centrifugation (800g for 10 min), and the supernatant was removed. The sample thus obtained was flash frozen in liquid nitrogen and stored at −80°C. Hi-C library preparation was performed as described previously (Belaghzal et al. 2017), except that ligated DNA size selection was omitted. 50 million fish liver cells were digested with *DpnII* at 37°C overnight. DNA ends were filled with biotin-14-dATP at 23°C for 4 hours. DNA was then ligated with T4 DNA ligase at 16°C overnight. Proteins were removed by treating ligated DNA with proteinase-K at 65°C overnight. Purified, proximally ligated molecules were sonicated to obtain an average fragment size of 200 bp. After DNA end repair, dA-tailing and biotin pull down, DNA molecules were ligated to Illumina TruSeq sequencing adapters at room temperature for 2 hours. Finally, the library was PCR-amplified and finalized following the Illumina TruSeq Nano DNA Sample Prep kit manual. Paired-end 50 bp sequencing was performed on a HiSeq4000.

The two Hi-C libraries from Connecticut and Georgia (the latter prepared by Dovetail) were mapped to the Atlantic silverside chromosome assembly using the *Distiller* pipeline (github.com/mirnylab/distiller-nf). Interaction matrices were binned at 50 and 100 kb resolution and intrinsic biases were removed using the Iterative Correction and Eigenvector decomposition (ICE) method (Imakaev et al. 2012). Large inversions (> 1 Mb) were identified by visual inspection of Hi-C maps as discontinuities that would be resolved when the corresponding section of the chromosomes were to be inverted (Dixon et al. 2018; Corbett-Detig et al. 2019). These discontinuities generate a distinct “butterfly pattern” with signals of more frequent Hi-C interactions where the projected coordinates of the breakpoints meet.

## Results

### Genome assembly and assessment of completeness

We obtained the best draft assembly (with the highest contiguity; N50 = 1.3 Mb) from the 10X data when we used 270 million reads as input to *Supernova*. Contiguity increased more than 2-fold with Dovetail Chicago data (scaffold N50 = 2.9 Mb) and more than 10-fold with Dovetail Hi-C data (scaffold N50 = 18.2 Mb). Summary statistics for each of the intermediate genome assemblies (10X, Dovetail Chicago, and Dovetail Hi-C) are presented in Table 1. The final assembly – including scaffolds longer than 1 kb only – was 620 Mb in total length. Overall, this assembly showed high contiguity, high completeness and a low proportion of gaps (Table 1). Analysis of the presence of BUSCO genes showed that only 5.9% of the Actinopterygii gene set were missing from the assembly. Although the number of missing genes did not decrease dramatically from the 10X assembly to the final assembly (from 6.6 to 5.9%), the addition of proximity ligation data (Chicago and Hi-C) increased the number of complete genes (from 88.1 to 89.6%) and decreased the number of duplicated (from 4.1 to 2.9%) and fragmented genes (from 5.3 to 4.5%). Contiguity did not come at the cost of increased gappiness, as stretches of N’s made up only 3% of the final assembly. The reduction of the assembly to its longest 27 scaffolds (‘chromosome assembly’-a 25% reduction in sequence) increased missing genes by only 3.1% and reduced duplicated genes to 1.9%. K-mer analyses based on raw data from the reference genome estimated a genome size of 554 Mb, 76 Mb shorter than the final assembly and 88 Mb longer than the chromosome assembly.

**Table 1.** Summary statistics for each of the intermediate and final assemblies produced.

|  | 10X | Dovetail Chicago | Dovetail Hi-C | Final assembly | Chromosome assembly* |
| --- | --- | --- | --- | --- | --- |
| Total length | 645.45 Mb | 647.32 Mb | 647.39 Mb | 620.04 Mb | 465.69 Mb |
| Longest Scaffold | 12,248,921 bp | 12,871,938 bp | 26,678,928 bp | 26,678,928 bp | 26,678,928 bp |
| Number of scaffolds | 99,541 | 80,990 | 80,312 | 42,220 | 27 |
| Number of scaffolds > 1kb | 61,451 | 42,898 | 42,220 | 42,220 | 27 |
| Contig N50 | 39.55 kb | 39.51 kb | 39.51 kb | 105.76 kb | 202.88 kb |
| Scaffold L50/N50 | 83/1.328 Mb | 42/2.936 Mb | 16/18.159 Mb | 15/18.199 Mb | 11/19.68 Mb |
| % gaps | 2.69% | 2.97% | 2.98% | 3.08% | 3.00% |
| BUSCOs**<br>(n=4584) | C:88.1%,<br>F:5.3%, M:6.6% | C:89.5%,<br>F:4.6%, M:5.9% | C:89.6%,<br>F:4.8%, M:5.6% | C:89.6%,<br>F:4.5%, M:5.9% | C:88.3%,<br>F:2.7%, M:9.0% |

\* The ‘chromosome assembly’ is the subset of scaffolds > 1 Mb from the ‘Final assembly’
\*\* [C=complete, F=fragmented, M=missing]

### Synteny with Medaka

The alignment of the 27 largest Atlantic silverside scaffolds to the medaka genome revealed a high degree of synteny conservation, especially considering the evolutionary distance between the two species. Each Atlantic silverside scaffold mapped mostly to only one medaka chromosome, and 22 of the 24 medaka chromosomes had matches with only one Atlantic silverside scaffold each (Fig. 1). Two medaka chromosomes, 1 and 24, had matches with three and two silverside scaffolds, respectively (Fig. 1). Based on these results, karyotype data confirming that the medaka and silverside have the same number of chromosomes (Uwa & Ojima 1981; Warkentine et al. 1987), and additional support from the Hi-C data from the Connecticut individual, we ordered and renamed the Atlantic silverside scaffolds according to the orthologous medaka chromosomes. We grouped the three and two scaffolds that mapped to medaka chromosomes 1 and 24, respectively, into one pseudo-chromosome each and renamed them accordingly. Although we did not observe large interchromosomal rearrangements in the alignment of the silverside and medaka genomes (Fig. 1), intrachromosomal rearrangements were common (Fig. 1; Fig. S1). The most conspicuous chromosomal rearrangements were large inversions, intrachromosomal translocations and duplications (Fig. 1; Fig. S1). On chromosomes 8, 11, 18 and 24, where large geographically differentiated haploblocks were identified among natural silverside populations, several translocations and inversions were evident, indicating poor intrachromosomal synteny (Fig. 1). This was also the case for most of the other chromosomes (Fig. S1).

**Figure 1.**
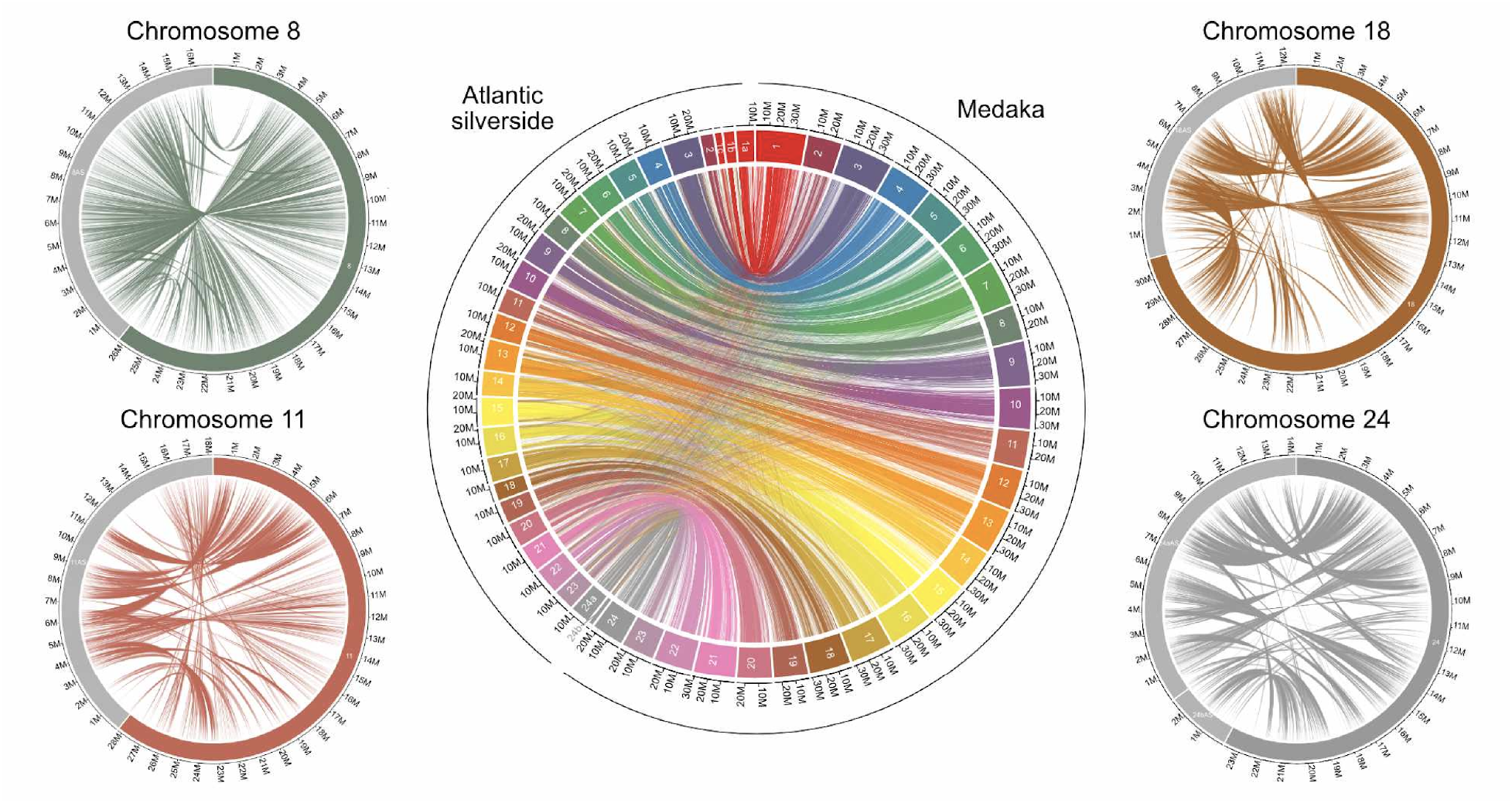
Circos plots showing synteny between the Atlantic silverside and medaka across all chromosomes in the middle and in the four chromosomes with large haploblocks on the sides. Chromosomes are color-coded consistently among plots and the colored portion of the smaller plots refer to the medaka sequences, while the grey portion to the Atlantic silverside sequences. Alignments shorter than 500 bp were excluded. Fig. S1 shows plots for the remaining chromosomes. Note that the consistently shorter length of the Atlantic silverside genome is consistent with a lower overall estimate of genome size (554 Mb based on k-mer analysis compared to the 700 Mb of the assembled medaka genome). The three and two scaffolds making up chromosomes 1 and 24, respectively, are represented separately here and denoted by small letters.

### Repeat and gene annotation

The identified repetitive elements made up 17.73% of the Atlantic silverside genome, in line with expectations based on fish species with similar genome sizes (Yuan et al. 2018). The biggest proportion of these repeats was made up of interspersed repeats (15.34% of the genome), while transposable elements constituted 8.83% of the genome overall (0.90% of SINEs, 2.79% of LINEs, 1.54% of LTR elements, and 3.60% of DNA elements). Our gene prediction pipeline identified a total of 21,644 protein coding genes, a number consistent with annotated gene counts in other fish species (Lehmann et al. 2019; Ozerov et al. 2018). Analysis in *Blast2GO* based on homology and *InterProScan2* resulted in functional annotation of 17,602 out of the 21,644 protein coding genes (81.3%; https://github.com/atigano/Menidia_menidia_genome/annotation/). Further, *InterProScan2* detected annotations (*Panther* or *PFAM*) for an additional 1,511 genes, for which no BLAST results were obtained.

### Sequence and structural standing variation

K-mer analyses based on raw data resulted in similar estimates of genome sizes and levels of heterozygosity in the two samples from Georgia and Connecticut: genome size estimates differed by 20 Mb (554 Mb and 535 Mb in the Georgia and Connecticut individual, respectively) and heterozygosity estimates differed by 0.09% (1.76% and 1.67% in Georgia and Connecticut, respectively). Direct estimates of heterozygosity, i.e. based on the number of called heterozygous sites in the genome, were slightly lower and differed by 0.14% between individuals (1.32% and 1.46% in Georgia and Connecticut, respectively). Together, these estimates concordantly indicate that standing sequence variation in this species is very high (Kajitani et al. 2014), with 1 in every ~66 bp being heterozygous within each individual. These heterozygosity estimates are higher than all comparable estimates reported for other fish species, though of similar magnitude to the European sardine and two eel species (Table 2). Heterozygosity varied substantially across the genome. Within each chromosome, heterozygosity was highest toward the edges of each chromosome, presumably in areas corresponding to telomeres, decreased towards the center in a U-shape fashion, and showed a deep dip in which the number of heterozygous sites approached zero, consistent with the location of putative centromeres (Fig. 2b). Additionally, the proportions of variable sites in coding regions was ~50% of whole genome level estimates (0.68% and 0.70% in Georgia and Connecticut, respectively). Swaths of low heterozygosity were particularly evident on chromosomes 18 and 24, two of four chromosomes with highly differentiated haploblocks (Fig. 2a,b).

**Table 2.**
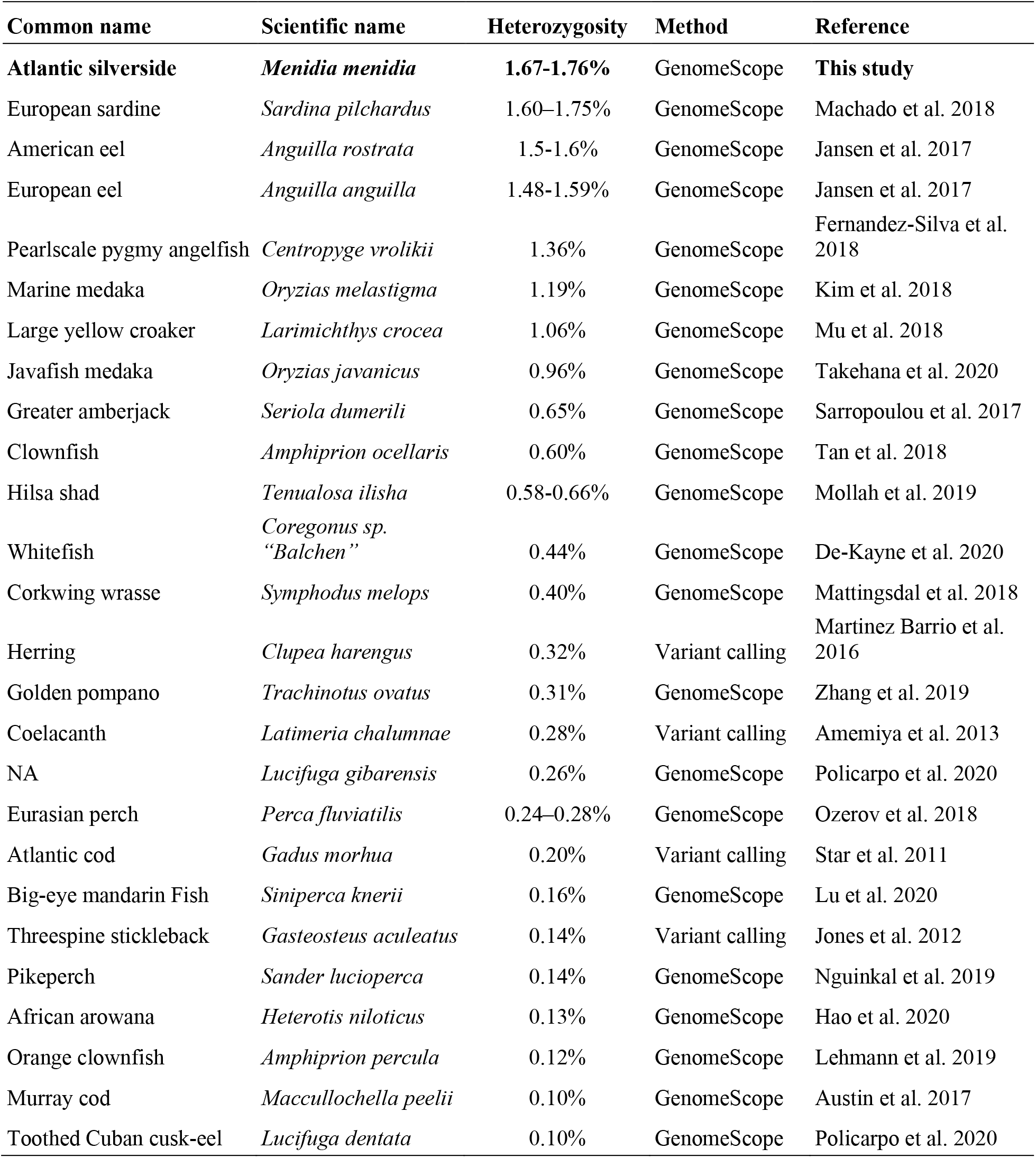
Examples of heterozygosity levels in single fish genomes, estimated either with GenomeScope from raw sequencing data or through direct calling of heterozygous sites.

**Table 3.** Summary of intraspecific structural variants identified in the Atlantic silverside, and their features.

| SV type | Number of variants | Size range (bp) | Sequence affected (kb) | % genome affected |
| --- | --- | --- | --- | --- |
| Insertions | 299 | 42-83 | 18 | <0.01% |
| Deletions | 3905 | 38-9,740,501 | 71,754 | 15% |
| Duplications | 34 | 110-150,263 | 479 | 0.1% |
| Inversions | 662 | 203-12,585,625 | 109,201 | 23% |

**Figure 2.**
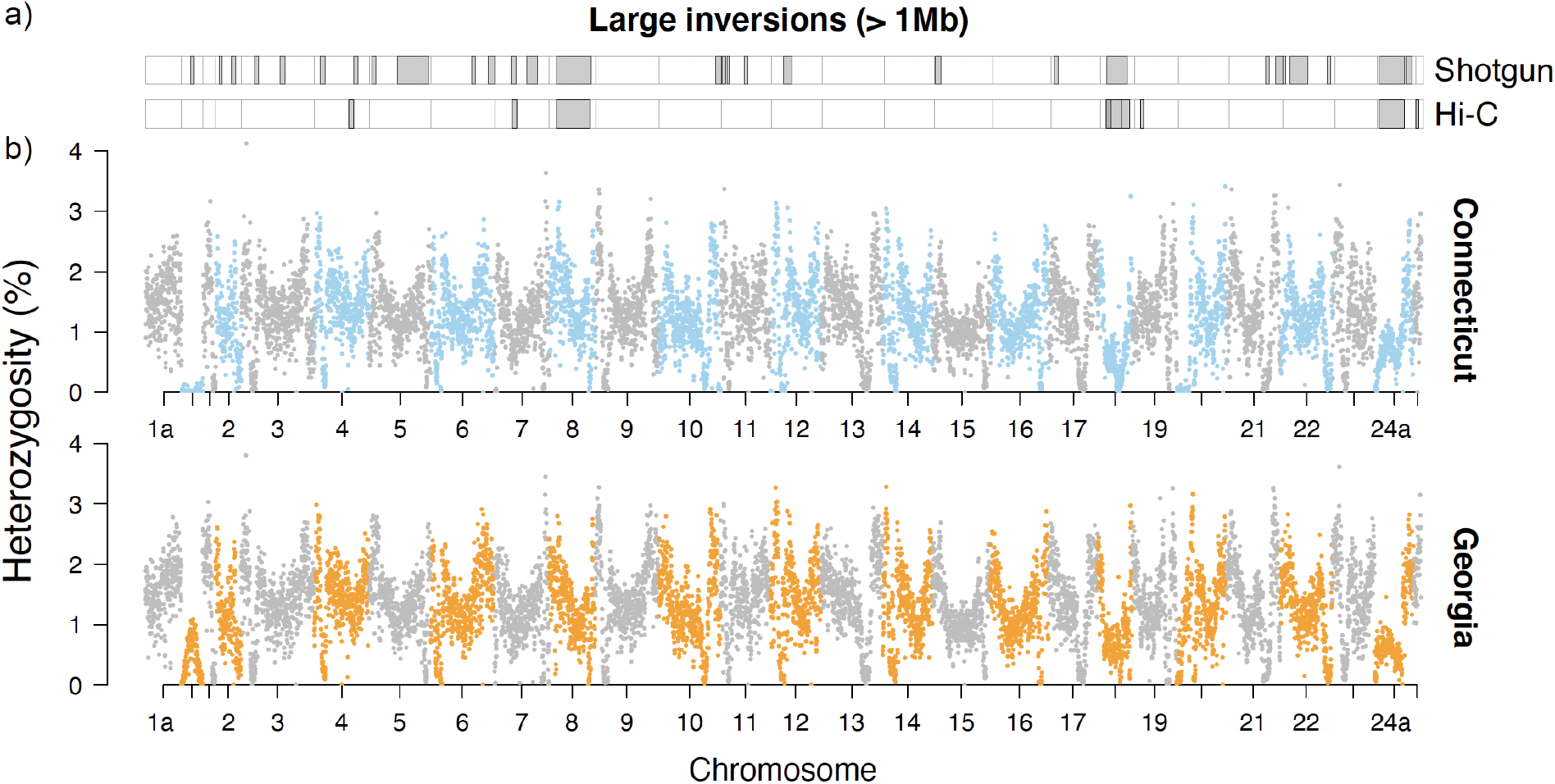
The genomic landscape of structural and sequence variation in Connecticut and Georgia. a) Panel showing large inversions (> 1 Mb) as identified from shotgun and Hi-C data from an individual from Connecticut mapped to the reference genome from Georgia. b) Manhattan plots showing the genomic landscape of variation in heterozygosity in 50 kb moving windows across single genomes from Connecticut and Georgia. The three and two scaffolds making up chromosomes 1 and 24, respectively, are represented separately here and denoted by small letters (e.g., 1a and 24a).

We identified a total of 4,900 SVs - including insertions, deletions, duplications and inversions (Supplementary File) - between the reference genome generated from Georgia samples and the re-sequenced individual from Connecticut. *Delly2* indicated that insertions were small (42-83 bp) and affected a negligible proportion of the genome, while deletions were larger and more abundant, covering 15% of the genome sequence. As an insertion in one genome corresponds to a deletion in the other genome depending on which individual is used as reference, the discrepancy between insertions and deletions is an artefact of mapping short-read sequences to a single reference, i.e. inserted sequences found only in Connecticut are not present in the reference and thus are not mapped. These results highlight the difficulties in identifying insertions and estimating their sizes from short reads. Our analysis detected a small number of duplications, covering only 0.1% of the genome. In contrast, we identified 662 inversions ranging from 203 bp to 12.6 Mb in size. In total, inversions affected 109 Mb, or 23%, of the reference genome sequence. Twenty-nine inversions were larger than 1 Mb, and five larger than 5 Mb (genomic locations in Fig. 2a and in Supplementary File). *Delly2* identified large inversions (> 1 Mb) on all four chromosomes with previously identified haploblocks. The largest inversion (~12 Mb) was identified on chromosome 8; chromosome 11 had two 1.2-Mb inversions that were 7 Mb apart; chromosome 18 had a 7.4 Mb inversion and chromosome 24 had two inversions, the first one spanning 9.4 Mb and followed by another one at a distance of 76 kb, spanning 2.3 Mb (Fig. 2a).

The independent Hi-C data from Connecticut (which was not used for genome scaffolding) supported a high degree of accuracy in the overall assembly into chromosomes, as indicated by the strong concentration of data points along the diagonal rather than elsewhere in the contact maps (Fig. 3). The contact maps also readily detected large-scale inversions (> 1 Mb) between the individual from Connecticut and the reference assembly from Georgia in three of the four chromosomes with haploblocks, i.e. 8, 18, and 24 (Fig. 3, Supplementary File). The missed detection of the inversions on chromosome 11 could either be due to their relatively smaller sizes, barely exceeding the detection threshold from Hi-C data, or because both inversion orientations segregate where the Connecticut individual used for Hi-C was sampled (Wilder et al. 2020). The breakpoints of the 12.6 and 9.4 Mb inversions on chromosomes 8 and 24, respectively, matched very closely those identified by *Delly2*, although the second 2.3 Mb inversion on chromosome 24 was not supported by Hi-C data (Figs. 2a, 3, Supplementary File). On chromosome 18, Hi-C data showed a complex series of nested and/or adjacent inversions spanning ~8.8 Mb in total, in contrast with the single inversion, and ~1.3 Mb shorter, identified by *Delly2* (Figs. 2a, 3, Supplementary File). Additional large inversions were detected from the Hi-C data on chromosomes 4, 7 and 19. Of these, the inversion on chromosome 19 was not identified from the analysis of shotgun data with *Delly2*, while those on chromosome 4 and 7 were, although with only one matching breakpoint for the inversion on chromosome 4 (Figs. 2a, 3, Supplementary File). Note that the identification of SVs from shotgun and Hi-C data were carried out by two different authors, and blindly from each other.

**Figure 3.**
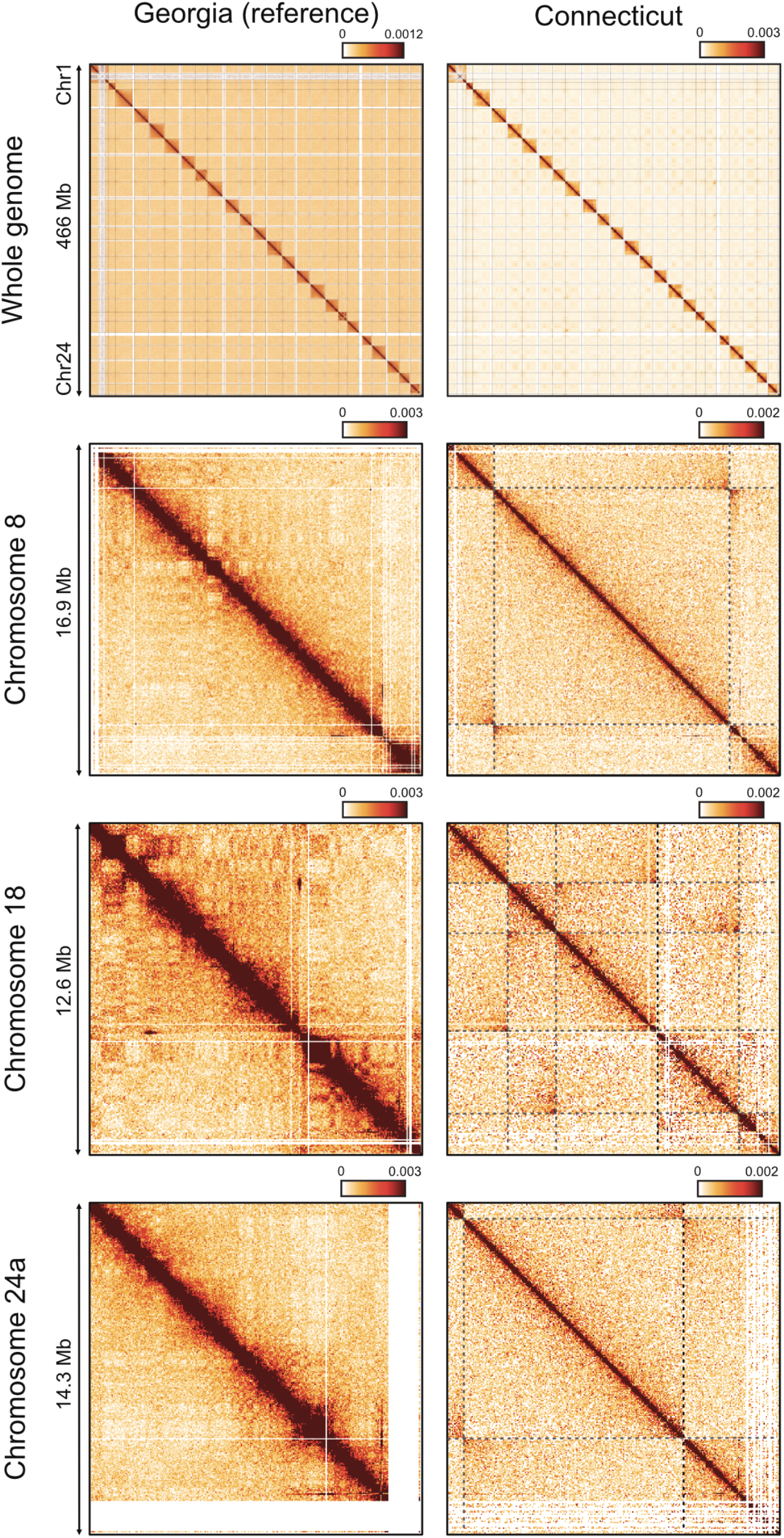
Hi-C contact maps of data mapped to the chromosome assembly from Georgia. Maps on the left show Hi-C data obtained from the same Georgia individual used to generate the reference assembly (mapped to self), maps on the right show data obtained from a Connecticut individual. Maps in the top panel show data for all the chromosomes binned in 100 kb sections. The three lower panels show data binned in 50 kb sections from each of the three chromosomes showing both large haploblocks in Wilder et al. (2020) and evidence for the presence of inversions from Hi-C data. Dark shades on the diagonal are indicative of high structural similarity between the reference and the Hi-C library analyzed. Dashed lines represent putative inversion breakpoints. The “butterfly pattern” of contacts observed at the point when the dashed lines meet is diagnostic of inversions.

## Discussion

We generated a chromosome-level assembly of the Atlantic silverside genome by integrating long-range information from synthetic long reads from 10X Genomics, *in vitro* proximity ligation data from Chicago libraries, and Hi-C proximity ligation data from whole cells. The resulting assembly had high contiguity and completeness. Based on karyotype information (Uwa & Ojima 1981; Warkentine et al. 1987), chromosome-level synteny with medaka, and Hi-C maps we reduced the 27 largest scaffolds to 24 putative chromosomes. This chromosome assembly is 88 Mb shorter than the genome size estimated through k-mer analysis, but has a lower number of duplicated genes, and only slightly fewer missing genes than the full assembly despite a substantial reduction in total sequence. If the proportion of complete genes in the chromosome assembly is, in fact, a good proxy for genome completeness, then the scaffolds that are not placed in chromosomes are mostly sequences that are repetitive, redundant, or that should fill gaps in the assembled chromosomes.

Heterozygosity within a sequenced individual can result in alternative alleles getting assembled into distinct scaffolds, even in genomes much less heterozygous than the Atlantic silverside (Kajitani et al. 2014; Tigano et al. 2018), so we expect some redundancy in our assembly. Considering the abundance of SVs between the two sequenced individuals, structural variation also may have contributed to the high number of smaller scaffolds not included in the chromosome assembly, as heterozygous SVs are notoriously hard to assemble (Huddleston et al. 2017). Nonetheless, the Atlantic silverside genome adds to the increasing number of high-quality fish reference genome assemblies, with the sixth highest contig N50 (202.88 kb) and the sixth highest proportion of the genome contained in chromosomes (84%, based on the genome size estimate from the k-mer analysis) compared to 27 other chromosome-level fish genome assemblies (Lehmann et al. 2019).

Patterns of synteny between the Atlantic silverside and the relatively distantly related medaka are consistent with comparisons among other teleost genomes up to hundreds of millions of years diverged: rearrangements are rare among chromosomes but common within (Amores et al. 2014; Rondeau et al. 2014; Miller et al. 2019; Pettersson et al. 2019). Consistent with this, anchoring Atlantic silverside transcriptome contigs on to medaka genome enabled the identification of four large haploblocks associated with fishery-induced selection in the lab and/or putative adaptive differences in the wild (Therkildsen et al. 2019; Wilder et al. 2020). However, the high degree of intrachromosomal rearrangements between the two species, and generally among teleosts, prevented an accurate characterization of the extent of these haploblocks and the analysis of structural variation. Differentiation between the northern and southern haplotypes seemed to extend across almost the entire length of three of the four chromosomes with haploblocks when data were oriented to medaka (Therkildsen et al. 2019; Wilder et al. 2020). However, the abundant intrachromosomal rearrangements between medaka and Atlantic silverside chromosomes (Fig. 1; Fig. S1), and the detection of large inversions in each of these four chromosomes (Figs. 2a,3) suggest that differentiation is concentrated in, and possibly maintained by, these inversions, which, albeit large, do not span whole chromosomes.

Our analysis of two genomes sequenced at high coverage suggested that levels of standing genetic variation, both sequence and structural, are extremely high in the Atlantic silverside. To our knowledge, our estimates of heterozygosity in a single individual are the highest reported for any fish species to date, including those with large census population sizes (Table 2). For example, heterozygosity, which is equivalent to nucleotide diversity (π) in one individual, in one single Atlantic silverside genome was higher than, or on par with, π estimates based on 43-50 individuals of Atlantic killifish, a species considered to have ‘extreme’ levels of genomic variation with π ranging from 0.011 to 0.016 (Reid et al. 2017, 2016). Compared to other vertebrates, genome heterozygosity in the Atlantic silverside was more than double the highest estimate reported for birds (0.7% in the thick-billed murre *Uria lomvia*; Tigano et al. 2018) and higher than the population-based 0.6-0.9% estimates in the rabbit (*Oryctolagus cuniculus*), one of the mammals with the highest genetic diversity (Carneiro et al. 2014). Among a collection of genome-wide π estimates - mostly population-based - across 103 animal, plant and fungal populations or species, only three insects and one sponge had π estimates higher than the Atlantic silverside (Robinson et al. 2016 and references therein). This unusually high level of standing sequence diversity is likely due to huge population sizes with estimated *Ne* exceeding 100 million individuals (Lou et al. 2018), and may underpin the remarkable degree of adaptive divergence and rapid responses to selection documented for the species.

Variation in π across the genome has been associated with variation in recombination rates, with higher diversity and recombination rates in smaller chromosomes and in proximity of telomeres in fish, mammals and birds (Ellegren 2010; Murray et al. 2017; Sardell et al. 2018; Tigano et al. 2020). In the Atlantic silverside, the decrease of heterozygosity from the ends towards the center of each chromosome is consistent with decreasing recombination rates as distance from the telomeres increases (Haenel et al. 2018; Sardell et al. 2018). However, in addition to this U-shape pattern, heterozygosity shows a dramatic, narrow dip in each chromosome far from the center of chromosomes, suggesting a strong centromere effect. Although striking differences exist between sexes and across taxa, recombination is generally reduced or suppressed around centromeres (Sardell & Kirkpatrick 2020). The Atlantic silverside karyotype, with only four metacentric and 20 non-metacentric chromosomes (i.e. submetacentric, subacrocentric, and acrocentric; Warkentine et al. 1987), further supports that these dips in heterozygosity are associated with centromeres, as the non-metacentric chromosomes enable the distinction between the effect of centromeres from the effect of distance from telomeres. In forthcoming work, linkage mapping will allow us to quantify the relative effects of centromeres and telomeres on local recombination rates and ascertain whether the recombination landscape is different between sexes.

We report a 50% reduction in heterozygosity in coding sequences compared to whole genome estimates, confirming the expectation that estimates based on exome data are not representative of whole-genome levels of standing variation. Even though the magnitude of the reduction in π within coding regions is similar to levels reported in the Atlantic killifish (Reid et al. 2017) and in the butterfly *Heliconius melpomene* (Martin et al. 2016), a substantially greater reduction is seen in the collared flycatcher (86%; Dutoit et al. 2017), suggesting that the distribution of diversity in a genome, including the difference between coding and non-coding sequence, is likely idiosyncratic to the population or species examined. Once again, a paucity of data from other species prevents us from making generalizations or identifying differences on the expected reduction in diversity in coding compared to non-coding regions across taxa, while at the same time it highlights the importance of estimating and reporting basic diversity statistics for whole genome assemblies.

We identified 4,900 structural variants that survived the stringent filters applied to maximize confidence in the identified SVs and to minimize the number of false positives due to genotyping one individual only. Our estimates are likely conservative when we consider that we filtered out all heterozygous SVs, that many SVs, particularly complex ones, are hard to identify or characterize (Chaisson et al. 2019), and that we analyzed only two genomes. Nonetheless, our analyses based on shotgun data show that SVs are abundant, affect a large proportion of the genome, with inversions covering up to 23% of the genome sequence, and range in size from small (< 50 bp) to longer than 10 Mb, with many of the largest inversions further supported by independent Hi-C data. Sunflower species of the genus *Helianthus* show a similar proportion of sequence covered by inversions (22%; Barb et al. 2014), although these were detected in comparisons between species (1.5 million years diverged) rather than within species. The few studies available on other species show that structural variation tends to affect a larger portion of the genome than single nucleotide polymorphisms (SNPs), but in proportions far lower than what we report here for the Atlantic silverside. For example, structural variation, including indels, duplication and inversions, covered three times more bases than SNPs did across six individuals of Australian snapper (*Chrysophrys auratus*; Catanach et al. 2019); short indels alone affected 4% of the genome of two individuals from the same population in the cactus mouse (*Peromyscus eremicus*; Tigano et al. 2020); inversions, duplications and deletions combined affected 3.6% of the genome across 20 individuals of *Tinema* stick insects (Lucek et al. 2019); and in cod (*Gadus morhua*) inversions covered ~7.7% of the genome (Wellenreuther & Bernatchez 2018 and references therein). Although levels of structural variation in the Atlantic silverside are extreme in comparison to these studies, a direct comparison with these and other species is hampered by a paucity of data and lack of common best practices for SVs genotyping (Mérot et al. 2020): differences in sampling, approaches, data types and filtering prevent comparisons similar to those made for standing sequence variation here and in other studies (Corbett-Detig et al. 2015; Robinson et al. 2016). Given the fast rate at which high-quality reference genomes are now generated, this will hopefully start to change.

The simple and affordable strategy we adopted only requires sequencing of a single additional shotgun library prepared from a second individual - possibly from a differentiated population to capture a broader representation of intraspecific variation - and could be easily applied in other studies to start describing variation in the prevalence and genome coverage of SVs across taxa. Here, an additional Hi-C library then allowed us to discover that the putative inversion on chromosome 18 was larger than indicated by the analysis of shotgun data and was actually constituted by a combination of two or more nested inversions. The apparent discrepancy between the breakpoints of the largest inversions identified using the two data types could reflect biological variation between the individuals analyzed. Alternatively, they may be caused by the different strengths and limitations of the underlying analytical approaches, including the fact that the identification of SVs was computational from shotgun data, while it was manually curated from Hi-C data. Although the analysis of only two individuals does not capture the full spectrum of intra- and inter-population variation, integrating different approaches has allowed us to identify a set of high-confidence SVs to be validated and genotyped in a larger number of individuals with lower coverage data (Mérot et al. 2020).

The joint analysis of sequence and structural variation reveals interesting features of the previously identified haploblocks. The chromosome-level assembly of the Atlantic silverside genome a) confirms that previously identified large haploblocks (Wilder et al. 2020) are associated with inversions and allows to measure their real extent; and b) highlights how genomic heterogeneity is multidimensional by revealing that even haploblocks showing similar patterns of differentiation can show vastly different patterns of genetic diversity. On chromosomes 18 and 24, large swaths of reduced heterozygosity (Fig. 2b) are associated with an inversion affecting the same area, which strongly indicates that the inversion promotes differentiation between genomes from Connecticut and Georgia in this region, likely through suppressed recombination. Of note, however, the segment of chromosome 24 preceding the inversion (0-722 kb) shows an even stronger reduction in heterozygosity than the adjacent inversion. While this additional reduction may be due to stronger recombination suppression in this area, the mechanism explaining this pattern remains to be investigated. In contrast, no reduction in diversity is associated with the inversion on chromosome 8 - the largest of them all (12.6 Mb) - or with the smaller inversions on chromosome 11. Such differences among haploblocks likely reflect idiosyncratic evolutionary histories and adaptive significance of the underlying inversions, whose investigation is now enabled by the chromosome-level genome assembly that we presented here. Hence, our analyses provide an empirical example of the importance of analyzing both sequence and structural variation to understand the mechanism underpinning the heterogeneous landscape of genomic diversity and differentiation.

Building on prior analysis based on in silico exome capture (Therkildsen & Palumbi 2017; Therkildsen et al. 2019; Therkildsen & Baumann 2020), this newly assembled reference genome provides an important resource for using the Atlantic silverside as a powerful model for investigating many outstanding questions in adaptation genomics, for example related to the abundance, distribution and adaptive value of structural variants; the relative role of coding and non-coding regions; the importance of sequence variation vs. structural variation in both human-induced evolution and local adaptation; and the demographic and evolutionary factors generating the genomic landscape of diversity and differentiation in this and other species.

## Acknowledgements

We would like to thank Hannes Baumann for rearing and collecting fish, Nicolas Lou for assistance with sampling, Harmony Borchardt-Wier for help in the lab, Leif Andersson for discussions on levels of heterozygosity in fish and other vertebrates, Peter Schweitzer at the Cornell Biotechnology Resource Center for advice on the 10X Genomics sample preparation and analysis, and Mark Daly at Dovetail Genomics for help with generating and interpreting the proximity ligation data. This work was funded by a National Science Foundation grant to N.O.T. (OCE-1756316) and the National Human Genome Research Institute (R01 HG003143 to J.D). J.D. is an investigator of the Howard Hughes Medical Institute.

## Author contributions

AT and NOT designed the study with input from JD; AJ performed the gene annotation; AW collected samples and performed lab work; AT, NOT, AW, YZ, AN and JD generated and analyzed the data; NOT and JD funded the project. AT wrote the paper with critical input from all authors.

## Data accessibility

The genome assembly and associated sequence data from Georgia and the raw data from the shotgun library from Connecticut will be available under ENA accession number #######. Scripts for the genome assembly and all other analyses can be found at https://github.com/atigano/Menidia_menidia_genome/.

