## Supplementary material for "Chromosome-level assembly of the Atlantic silverside genome reveals extreme levels of sequence diversity and structural genetic variation": Figure S1

**Supplementary Figure 1**. Circos plots showing synteny between the Atlantic silverside and medaka for each chromosome. Chromosomes are color-coded consistently with Figure 1 in the main text. The colored portion of the plots denotes medaka sequences, while the grey portion denotes Atlantic silverside sequences (note that the consistently shorter length of the Atlantic silverside genome is consistent with a lower overall estimate of genome size (554 Mb based on k-mer analysis compared to the 700 Mb of the assembled medaka genome). Alignments shorter than 500 bp were excluded. First page includes chromosomes 1-12, second page includes chromosomes 13-24.

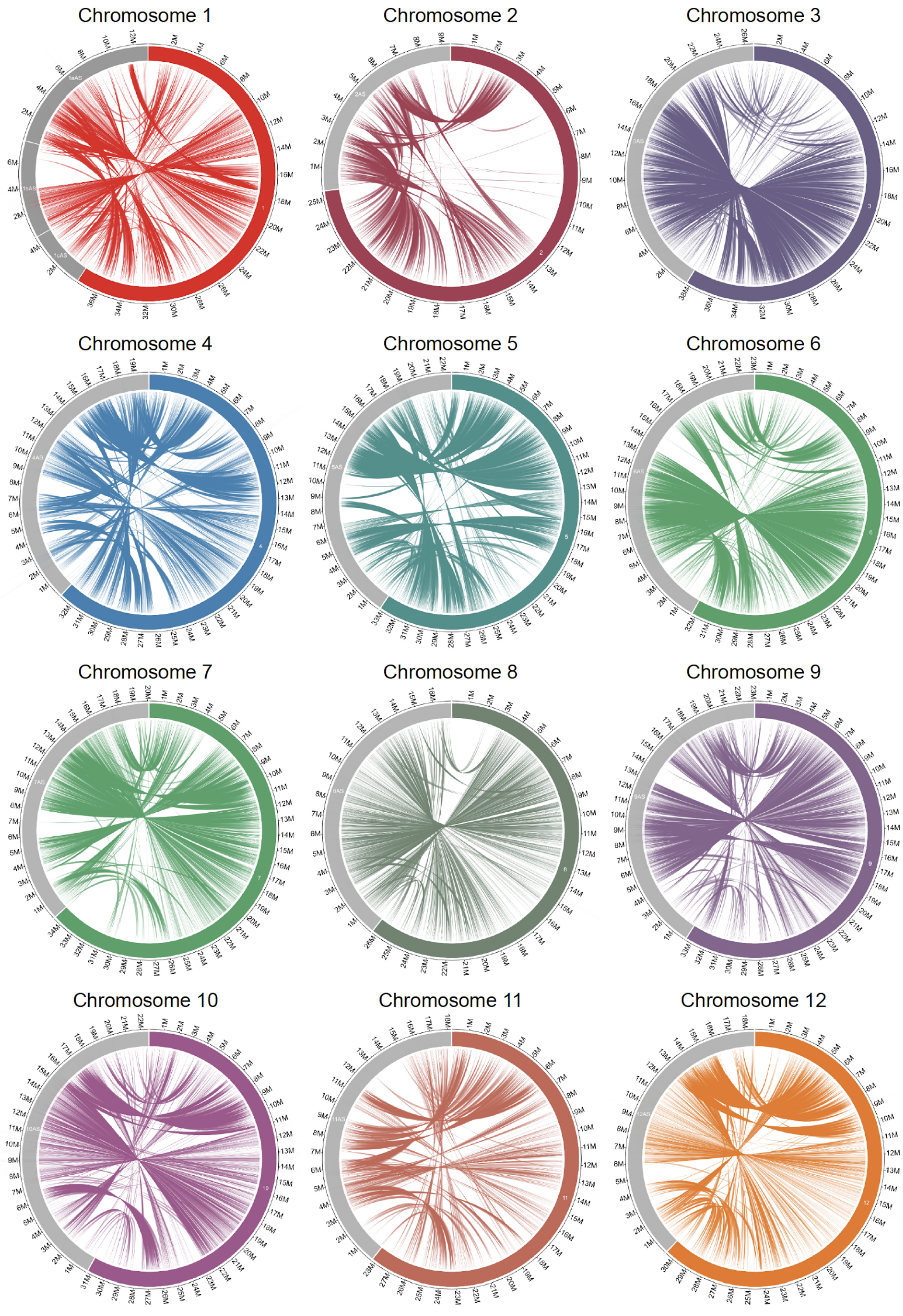

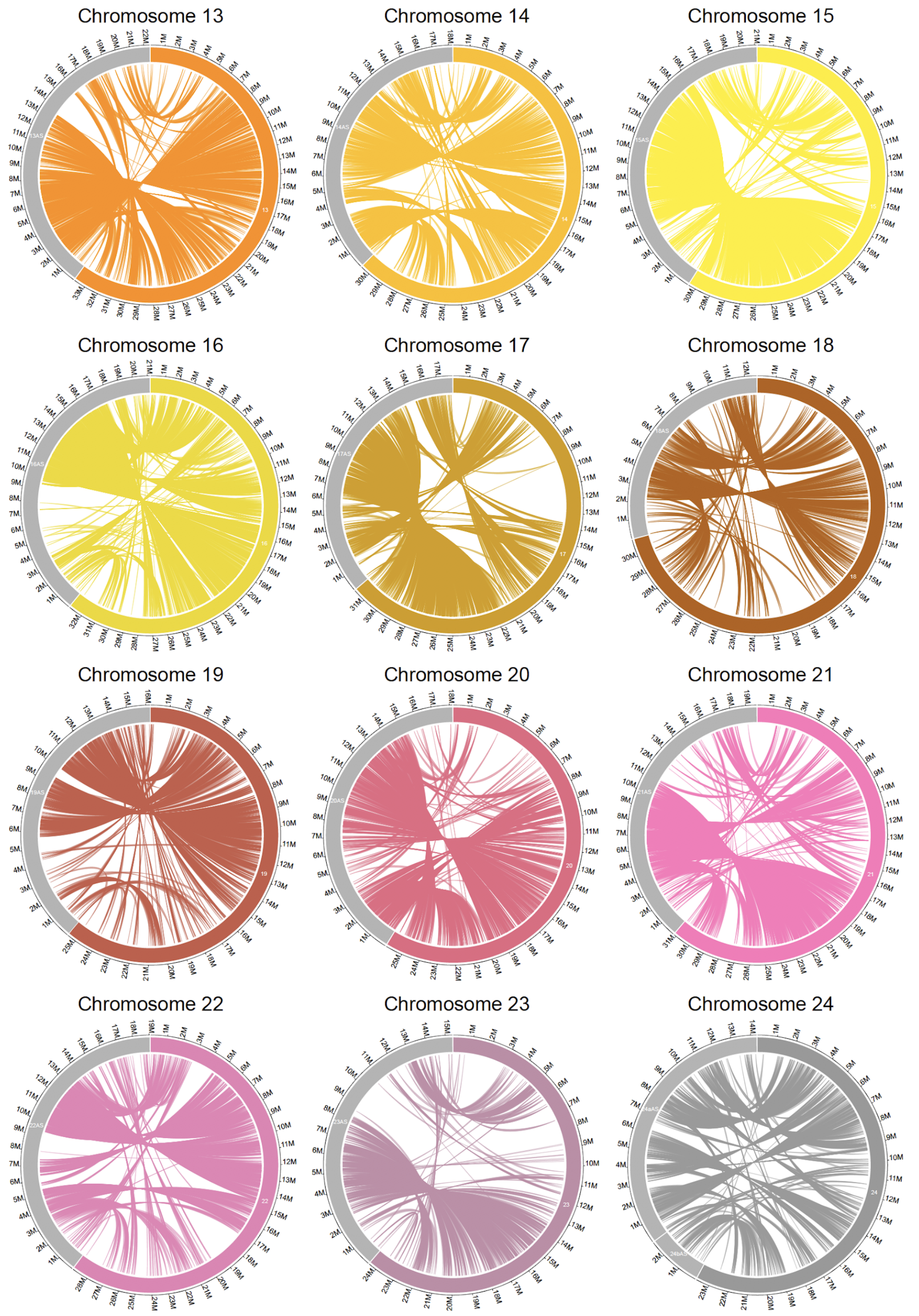
Chromosome 13
Chromosome 14
Chromosome 15
Chromosome 16
Chromosome 17
Chromosome 18
Chromosome 19
Chromosome 20
Chromosome 21
Chromosome 22
Chromosome 23
Chromosome 24
